# Urban tree diversity fosters bird insectivory despite a loss in bird diversity with urbanization

**DOI:** 10.1101/2024.11.15.623789

**Authors:** Laura Schillé, Alain Paquette, Gabriel Marcotte, Hugo Ouellet, Swane Cobus, Luc Barbaro, Bastien Castagneyrol

## Abstract

Urbanization is one of the main drivers of biotic homogenization in bird communities worldwide. Yet, only a few studies have addressed its functional consequences on the top-down control birds exert on insect herbivores. We hypothesized that their inconsistent results reflect the overlooked heterogeneity of the urban habitat for birds, and in particular the distribution and diversity of urban trees.

We monitored tree diversity, bird diversity, avian predation attempts on artificial prey, and the effect of bird exclusion on insect herbivory in 97 trees distributed among 24 urban experimental plots in the city of Montreal, Canada. We characterized urbanization levels through a combination of variables related to tree density, impervious surfaces, anthropic noise, and human population density.

Bird diversity decreased with increasing urbanization, whereas the frequency of generalist synurbic species increased. We found no significant relationship between predation and urbanization or between predation and bird diversity. However, tree diversity was positively correlated with predation attempts on artificial prey, irrespective of bird diversity.

We revealed a mismatch between the effects of urbanization on bird diversity and on the regulation service and unraveled the functional importance of tree diversity in shaping the avian predation function in urban ecosystems. Our study advocates for the consideration of intra-urban heterogeneity in the investigation of trophic cascades within cities.

## 1. Introduction

Populations of insectivorous birds are declining worldwide (Bowler et al., 2019; Rosenberg et al., 2019). Causes are numerous and likely interacting. They include a sharp decline in the abundance of insect herbivores that are a main source of food for insectivorous birds (Bowler et al., 2019), the intensification of anthropic pressure on ecosystems (Devictor et al., 2007), and climate change (Both et al., 2006). Yet, insectivorous birds provide humans with essential regulation services by reducing insect herbivore pressure on plants through top-down trophic cascades (Nyffeler et al., 2018). The factors influencing the strength of bird-induced trophic cascades have been well-documented in natural and cultivated ecosystems (Harris et al., 2020; Jactel et al., 2021). However, whether and to which extent knowledge gained in these ecosystems can be transferred to urban environments remains uncertain (Frey et al., 2018; Kozlov et al., 2017; Long & Frank, 2020).

Both the abundance (Roels et al., 2018), activity (Maas et al., 2015), and diversity (Nell et al., 2018) of insectivorous birds are well-documented predictors of the top-down control they exert on insect herbivores. Insectivory increases with the abundance and foraging activity of insectivorous birds, particularly during the chick feeding period (Naef-Daenzer & Keller, 1999). It is also a period during which non-insectivorous species may include insects in their diet to feed their nestlings. Theory predicts that predation pressure increases with increasing diversity of insectivorous birds, as a result of functional complementarity among species within the community (Philpott et al., 2009). Several studies consistently reported positive relationships between the functional diversity of insectivorous birds and the regulation function (Barbaro et al., 2014; Peña et al., 2023). In the same way, any environmental factor influencing the distribution, diversity, or composition of insectivorous bird communities is likely to also affect the trophic interactions they initiate, but the strength and direction of these effects are elusive (McDonnell & Hahs, 2015).

Urbanization alters the composition of bird communities (McKinney, 2006) as well as bird behavior (Fuller et al., 2007). The scarcity of food and nesting opportunities generally reduces the taxonomic and functional diversity of birds (La Sorte et al., 2018) making urban bird communities dominated by habitat and diet generalists, as well as poorly mobile, generally resident species (Lakatos et al., 2022). In addition, the distribution and activity of birds are disrupted by anthropogenic nuisances such as noise (Slabbekoorn & Ripmeester, 2008), light pollution (Aulsebrook et al., 2020), direct disturbances by humans (Price, 2008) or by their domestic carnivores (Baker et al., 2005). All those disturbances are also likely to modify the ability of bird communities to regulate herbivorous arthropods. However, how urbanization affects predation is still poorly documented and contradictory. Some studies showed a decrease of herbivore regulation in urban ecosystems, when compared to non-urban ones (e.g., Turrini et al., 2016), whereas others highlighted a stronger top-down control of herbivores in urban areas (e.g., Faeth et al., 2005). The same contradictions occur for the arthropod herbivory in urban versus rural environments (Kozlov et al., 2017; Raupp et al., 2010). A possible explanation for these inconsistencies might be the focus on city-to-countryside gradients, and the lack of consideration for intra-urban heterogeneity.

Indeed, cities are heterogeneous environments in which the abundance and distribution of trees jointly contribute to shaping biodiversity. Locally, tree diversity has the potential to alleviate the adverse effect of urbanization on avian communities (Da Silva et al., 2021) and on the biocontrol they exert on herbivores (Stemmelen et al., 2020). The diversity of insect herbivores and their natural enemies generally correlates positively with tree diversity (Castagneyrol & Jactel, 2012) as a result of increased habitat complexity and food resources for insectivorous birds (Barbaro et al., 2019). As stated by the natural enemies’ hypothesis (Root, 1973), the regulation function provided by birds is thus expected to increase with the diversity of trees. Efforts to restore tree biodiversity in urban areas could therefore allow for the maintenance of higher bird diversity and their biotic interactions, a key issue for managing urban ecosystems.

In this study, we investigated the interacting effects of urbanization and tree diversity on bird diversity and insectivory, to infer their consequences on tritrophic interactions between birds, insect herbivores, and trees. To document the ecological functions provided by trees and their diversity in urban environments, we measured tree diversity, insect herbivory, insectivorous bird diversity, and the regulation function provided by birds in 24 urban plots distributed along a gradient of urbanization within the city of Montreal, Canada. We predicted the following : (i) urbanization reduces insectivorous bird diversity and predation and thereby the strength of tri-trophic cascade, leading to increased insect herbivory on trees; (ii) local tree diversity is positively associated with insectivorous bird diversity and predation and (iii) local tree diversity mitigates the effect of urbanization.

## 2. Material and Methods

### 2.1. Study area

The study was conducted in the city of Montreal, Canada, which has a population of approximately 1.8 million inhabitants. It was conducted during two weeks across May and June 2023 (from May 22 to June 5). The average temperatures for these two months were 15.9°C and 21.5°C, respectively (Pierre Elliott Trudeau airport weather station). We used a pre-established network of 24 urban plots called the Urban Observatory of Montreal (https://paqlab.uqam.ca/tags.php?id=45&lang=fr, Fig.1) selected to cover the largest range of two gradients: (i) tree density, approximated through the percentage of canopy cover, (ii) human population density, used here as proxy for human disturbance in bird communities.

**Figure 1.**
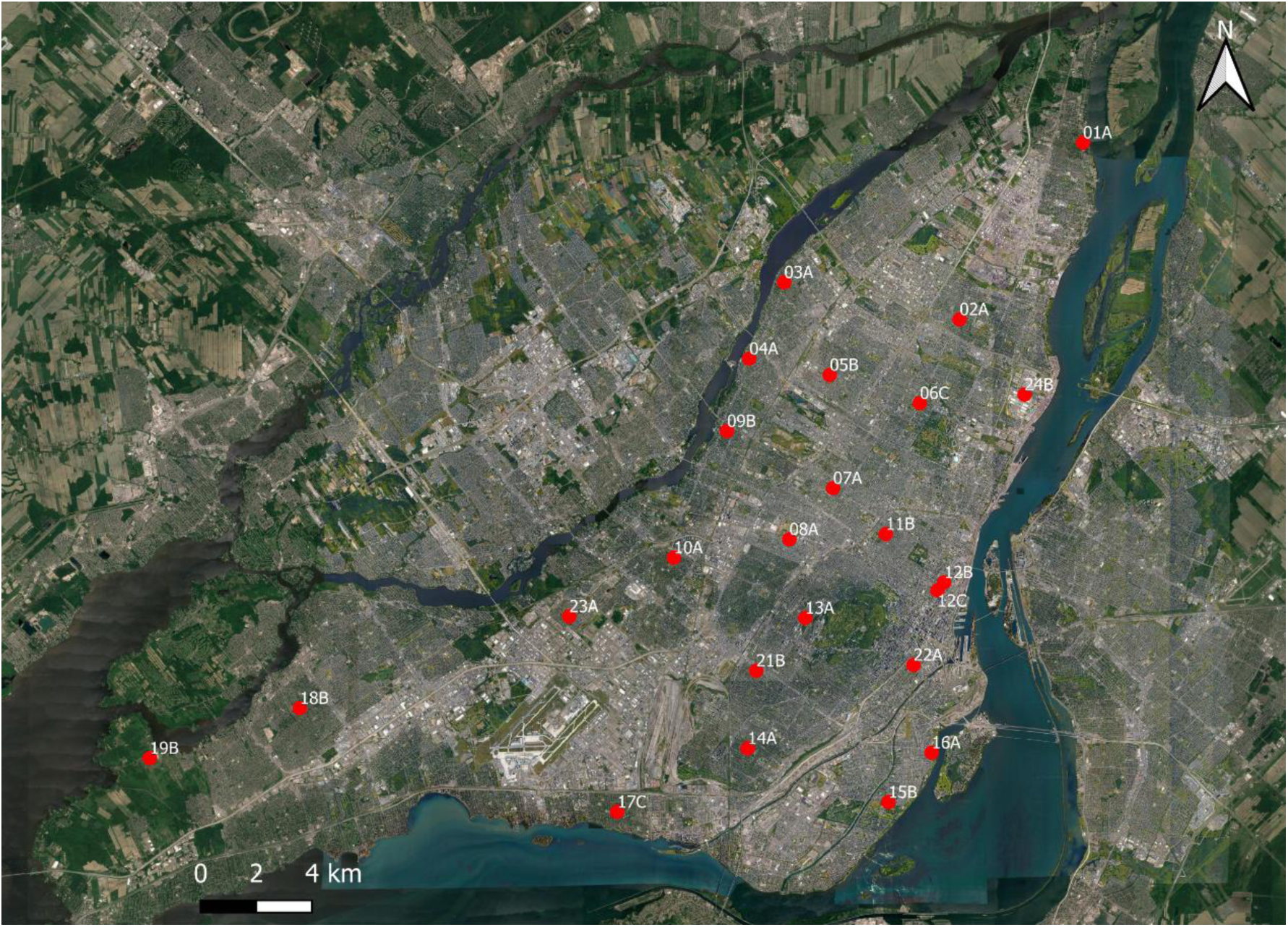
Locations of the 24 sampled urban plots in the city of Montreal, Canada.

### 2.2. Focal trees and neighborhood

We selected four deciduous trees (henceforth, focal trees) in each urban plot, except for one plot where we selected five trees because we were concerned that the experimental material at this site was particularly at risk of vandalism. We chose focal trees among the ten most abundant species in Montreal, namely *Acer platanoides* (non-native), *Acer saccharinum* (native), *Acer negundo* (native), *Gleditsia triacanthos* (non-native), *Tilia cordata* (non-native), *Tilia americana* (native), *Prunus* spp. (cultivars), *Ulmus americana* (native), *Ulmus pumila* (non-native), and four *Ulmus* x *hybrids* (cultivars). We ensured that focal trees met the following criteria: be within a 100m radius of the plot center (to link with tree richness measured over a 100m radius), have easily accessible branches using a small ladder, and be located in the public domain. We further ensured that each focal tree species was repeated at least three times in three different urban plots. This resulted in the selection of 97 focal trees (Table SA).

Tree diversity around each focal tree was characterized through the prior identification of all trees in both the public and private domains within a 200m radius around the plot center. Without authorization from the property owners, the trees were identified with binoculars from the street. We used tree species richness in a 20m radius circular buffer to characterize the neighborhood of each focal tree. We could not extend the calculation of neighbor tree species richness beyond 20m to focal trees as some of them were located too close to the edge of the plots. Eventually, we average neighboring tree species richness at the plot level. We were unable to calculate a neighborhood tree diversity index because we had no information on the diameter at breast height for 35% of the neighboring trees.

### 2.3. Insectivorous bird species richness and functional diversity

We used passive acoustic monitoring to identify all vocalizing bird species and characterize bird communities within the plot, at the species level. We selected a central tree or hedge in each urban plot to conceal an AudioMoth device (Hill et al., 2018). The AudioMoth was parameterized to record audible sounds for 30 minutes every hour. Automated recording began on the day we installed the plasticine caterpillars in trees, for eight consecutive days.

For every urban plot, we only sub-sampled the 30-minute recordings corresponding to the morning chorus of songbirds (from 30 minutes before sunrise to 3 hours and 30 minutes after sunrise). We only retained the subsamples recorded on Tuesday, Thursday, Saturday and Sunday to balance out the impact of human activity between weekends and weekdays. These 30-minutes recordings were then divided into three 10-minutes segments. Each sound sample underwent analysis through the ’seewave’ library in the R environment (Sueur et al., 2008), and any sample containing low-frequency noise typical of human activity, rain, or wind were excluded based on their spectrogram appearance. This process yielded a total of 96 10-minutes recordings (4 recordings per urban plot).

All recordings were listened to by a single expert ornithologist (H.O.), who identified all species heard, noting the precise time intervals during which each species vocalized. For each vocalization interval, the expert recorded the species present as well as the number of vocalizations emitted by each of them. The number of occurrences for each species was then summed across the recording, and an average of the occurrences per species from the four recordings at each site was calculated. It should be noted that this process provides a vocalization occurrence of each species and, therefore, a form of activity of that species, but it is not a reliable measure of bird species abundance (Schillé, Valdés-Correcher, et al., 2024). We focused exclusively on functionally insectivorous birds, defined as species that meet the following criteria: being totally or partially insectivorous during the breeding period or feeding their chicks with insects, foraging primarily in trees, and using lower branches of trees to find their prey (see Fig. 3 for the characterization of bird species on their diet).

Using the presence/absence Plot × Species matrix, we computed the species richness of the communities in each urban plot, and we used the vocalization occurrence Plot × Species matrix to calculate a vocalization-based Shannon index (hereafter referred to as Shannon diversity). Additionally, we integrated morphological, reproductive and behavioral traits (Cornell Laboratory of Ornithology, 2022) to compute the vocalization-based functional dispersion index (hereafter referred to as FDis), an index representing species dispersion in trait space, weighted by their relative vocalization occurrence. This index was calculated for each urban plot using the dbFD function of the FD library (Laliberté et al., 2014) in the R environment.

We calculated the Bioacoustic Index (BI) in each recording as a proxy for overall bird activity. The BI measures the area under the sound spectrum curve, above a certain intensity threshold, and within a specific frequency range (Boelman et al., 2007). Previous studies have shown that this index is positively correlated with bird abundance (Boelman et al., 2007) and their predation function (Schillé et al., 2024). We consider this measure particularly informative because it more directly reflects the proportion of sound signal produced by vocalizing species, rather than merely counting the number of vocalizations. To calculate this index, we first segmented the entire set of recordings over eight days into one-minute intervals. We then used a wrap-up function developed by A. Gasc, available on GitHub (https://github.com/agasc/Soundscape-analysis-with-R), to determine the BI for each minute of every recording at each site. Eventually, we calculated the median BI for each day and averaged these median values across all days for each urban plot.

### 2.4. Urban habitat variables

We integrated information from eight variables representing human activities and habitat characteristics to refine the definition of the urbanization gradients encompassing the 24 urban plots (Table 1). When the data were accurate enough, we used an aggregation level of 200m around the focal trees. Indeed, preliminary analyses revealed that the results were qualitatively identical using buffers of 100, 200, or 500m, as the metrics used were highly correlated at these scales. We chose a buffer size of 200m because several common insectivorous birds in the city of Montreal have vital ranges not exceeding 200m. This is the case, for example, for the Black-capped Chickadee, whose vital range can go from 1.5 ha to 5.3 ha (equivalent to a circle with a radius of 130m) (Québec center of expertise in environmental analysis, 2005) or Warblers with average vital ranges of 2 ha (Cornell Laboratory of Ornithology, 2022).

**Table 1.** Summary of the variables used to describe urbanization, their definition, their sources, and their aggregation level.

| Used variable | Variable definition | Data source | Aggregation level |
| --- | --- | --- | --- |
| Imperviousness | Percentage of the combined surface area of low and high mineral surfaces | Normalized Difference Vegetation Index (NDVI) and city's digital elevation model (Communauté métropolitaine de Montréal, 2017) | 200m around focal trees |
| Building cover | Percentage of high mineral surface area | NDVI and city's digital elevation model (Communauté métropolitaine de Montréal, 2017) | 200m around focal trees |
| Canopy cover | Percentage of high vegetation surface area | NDVI and city's digital elevation model (Communauté métropolitaine de Montréal, 2017) | 200m around focal trees |
| Population density | Average household size from the 2016 census | Population census data (Government of Canada 2016) | 200m around focal trees |
| Light pollution | Night-time radiance | Satellite VIIRS image (National oceanic and atmospheric administration, 2023) | Plot level |
| Sound intensity |  | Measured with a sound level meter in the field | Four measuring points per focal tree |
| Habitat connectivity | Number of high vegetation patches | Derived from NDVI and city's digital elevation model (Communauté métropolitaine de Montréal, 2017) | 200m around focal trees |
| NDSI index | Biophony to anthropophony ratio | Calculated from passive acoustic field recordings | Plot level |

Specifically, we used the metropolitan canopy index raster layer and QGIS software (version 3.16 Hannover) to extract the percentage of imperviousness, of building density and of canopy coverage at the tree level. We also calculated habitat connectivity at the tree level. The values were then averaged at the plot to standardize the scale across different urbanization metrics.

Population density was extracted from a vector layer of the 2016 census. We calculated an average estimate of population density around the target trees and then averaged the values at the plot level.

We used the VIIRS night-time radiance raster layer, an annual cloud-free composite with a 450m pixel resolution, to extract light pollution at plot level.

The sound intensity was measured using a digital sound level meter (*TROTEC, SL400, measurement range from 30 to 130 dB, slow sampling rate, i.e., one measurement per second*) for 5 minutes each time we visited focal trees to install or remove the material or collect samples at one-week interval or more. This resulted in 16 measurements per urban plot (4 dates × 4 trees), which were taken on different weekdays and times during the day, with no particular sequence to avoid confounding effects. The measurement device was consistently positioned with its back to the tree, pointing toward the nearest road, and placed on a tripod at 1m above the ground with a 60° inclination relative to it. We averaged the data across sampling dates and trees to provide a synthetic estimate of anthropogenic noise in each urban plot.

Finally, we characterized the soundscape of the urban plots using the recordings made with the Audiomoths. To do so, we first cut each recording into 1-minute chunks. We then calculated the ratio between biophony and anthropophony using the Normalized Difference Sound Index (NDSI) thanks to ’soundecology’ library in the R environment (Villanueva-Rivera & Pijanowski, 2018). We then calculated the median of NDSIs per day and then averaged median values across days for each urban plot individually.

### 2.5. Bird predation attempts

We assessed the activity of insectivorous birds by counting predation attempts (as revealed by bill marks) on plasticine caterpillars (Low et al., 2014). We made 3 × 0.5 cm caterpillar-shaped cylinders of green plasticine (brand *Staedtler*, model 8421-5). We exposed 20 plasticine caterpillars per tree, on twigs with a 0.3 mm diameter metallic wire. We attached five plasticine caterpillars to four branches facing opposite directions, about 2.5 m from the ground. We installed them five weeks after the average budburst date for all the selected species, thus synchronizing the study with local tree phenology. They were exposed for 15 days and were removed one week before we collected leaves for herbivory measurements. Predation marks on the plasticine caterpillars were only assessed once, after their removal, directly in the field by a single trained observer (L.S.) who examined each caterpillar, checking for the presence of bill marks on the clay surface. When there were doubts about the type of attack, the caterpillars were brought back to the lab for a second check and to seek advice from colleagues. We approximated bird predation attempts through the number of plasticine caterpillars presenting at least one evidence of bird attack.

### 2.6. Bird exclusion

To assess the strength of the trophic cascade, we prevented insectivorous birds from foraging on one branch per focal tree. To this end, we installed a bird exclusion net around a 1m long branch in April 2023, a few days before the expected budburst. Nets were made of green plastic material and had a mesh size of 15 × 15mm to prevent bird access while allowing the passage of herbivorous arthropods). The net was adjusted weekly throughout the budburst process to ensure that leaves remained enclosed within the net.

### 2.7. Insect herbivory

We compared insect herbivory in the bird exclusion treatment with insect herbivory on control branches that were adjacent to those with the exclusion treatment. Specifically, we haphazardly collected 30 leaves in each treatment, totaling 60 leaves per focal tree. We collected leaves eight weeks after budburst, which corresponds to the peak of activity of insect herbivores and to the period during which we assessed bird diversity and activity. We oven-dried the leaves for 48 hours at 45°C and haphazardly selected 20 leaves (out of 30) per treatment and per tree to be processed for further analyses.

Herbivory was visually assessed by a single experimenter (G.M.) and was defined as the percentage of the leaf surface removed or impacted by insect herbivores. The leaves were assigned to 11 different classes based on the amount of damage: 0%, [1%-5%], [6%-10%], [11%-15%], [16%-20%], [21%-25%], [26%-30%], [31%-40%], [41%-50%], [51%-75%] and >75%. We then used the median values of these classes to calculate an average herbivory rate across the 20 leaves per treatment and per tree. We defined the herbivory regulation function provided by birds as the difference between the herbivory on the netted branch and the herbivory on the control branch.

### 2.8. Statistical analyses

We first summarized the information about urbanization by running a Principal Component Analysis (PCA) on the eight urbanization-related variables. By doing so, we intended to provide a synthetic and reliable description of the degree of urbanization in each urban plot. We extracted the coordinates of the projection of plots onto the first two PCA axes, which together explained 71% of the variance in urban plot descriptors (see *Results*). We used PCA coordinates as proxies for urbanization intensity (PC1) and for urban naturalness (PC2) in further analyses (see below).

We tested the effects of urbanization (PC1 and PC2) and neighborhood tree species richness on insectivorous birds (species richness, Shannon diversity, FDis, and BI), their activity (predation attempts on plasticine caterpillars), and the regulation function they provide using separate models (Table SB). We approximated the regulation function provided by insectivorous birds through the difference between insect herbivory on branches from which birds were excluded and insect herbivory on control branches.

We analyzed bird-related variables with data aggregated at the plot level using Linear Models (LMs) or Generalized Linear Models (GLMs). We used GLM with Poisson error distribution to model count data (bird species richness), LMs with Gaussian error distribution to model bird diversity (Shannon diversity and FDis), and LM with Gaussian error distribution with a prior square root transformation of the response variable for the Bioacoustic Index (BI) to meet the LM assumptions. In each model, predictors were PC1 and PC2 coordinates along with neighborhood tree species richness and every two-way interaction.

We used a Generalized Linear Mixed Model (GLMM) with binomial error distribution to analyze predation attempts (i.e., number of attacked vs. number of intact plasticine caterpillars), and an LMM with Gaussian error distribution to analyze herbivory regulation function. As for the bird-related models, predictors were PC1 and PC2 coordinates along with tree species richness and every two-way interaction. Because we predicted that the effect of urbanization and tree diversity on the tri-trophic cascade would be mediated by changes in insectivorous bird diversity or activity, we included insectivorous bird species richness or BI as an additional predictor. We excluded models that contained both variables simultaneously to ensure a clearer interpretation of their individual effects. Urban plots and tree species were considered as crossed random effects in GLMMs. Preliminary analyses indicated that the native or exotic status of the focal tree did not account for variability in predation attempts or in regulation function provided by insectivorous birds. For the sake of parsimony, we did not consider this variable in further analyses.

We scaled and centered every continuous predictor other than PC1 and PC2 before modeling to facilitate comparisons of their effect sizes. We made sure that none of the explanatory variables were strongly correlated by examining the variance inflation factors (VIF) (all VIFs < 5, a common cutoff value used to check for multicollinearity issues).

For each response variable, we ran the full model as well as every model nested within the full model and then used Akaike’s Information Criterion corrected for small sample size (AICc) to identify the most parsimonious models best fitting the data. Models within 2 ΔAICc units of the best model (i.e., the model with the lowest AICc) are generally considered as equivalent in their ability to fit the data the best. We computed AICc weights for each model (*wi*), where *wi* is interpreted as the probability of a given model being the best model among the set of candidate models. Eventually, we calculated the relative variable importance (RVI) as the sum of *wi* of every model including this variable, which corresponds to the probability a variable is included in the best model.

When several models competed with the best model (i.e., when multiple models were such that their ΔAICc < 2), we applied a procedure of multimodel inference, building a consensus model including the variables in the set of best models. We then averaged their effect sizes across all the models in the set of competing models, using the variable weight as a weighting parameter (i.e., model averaging). We considered that a given predictor had a statistically significant effect on the response variable when its confidence interval excluded zero.

We ran all analyses in the R language environment (R Core Team, 2021) with libraries “MuMIn” v.1.43.17 (Bartoń, 2020), “lme4” v. 1.1.27.1 (Bates et al., 2015).

## 3. Results

### 3.1. Main urbanization drivers

The first two dimensions of the PCA explained 71% of the variance in urbanization-related variables (Fig. 2). PC1 was positively associated with sound intensity, human population density, building density, and impervious surfaces, with positive values representing urban plots with higher urbanization intensity. PC2 was positively associated with canopy cover and connectivity among forest patches, and negatively associated with NDSI and light intensity. It therefore represented a gradient of urban naturalness whereby positive values represent greener urban plots.

**Figure 2.**
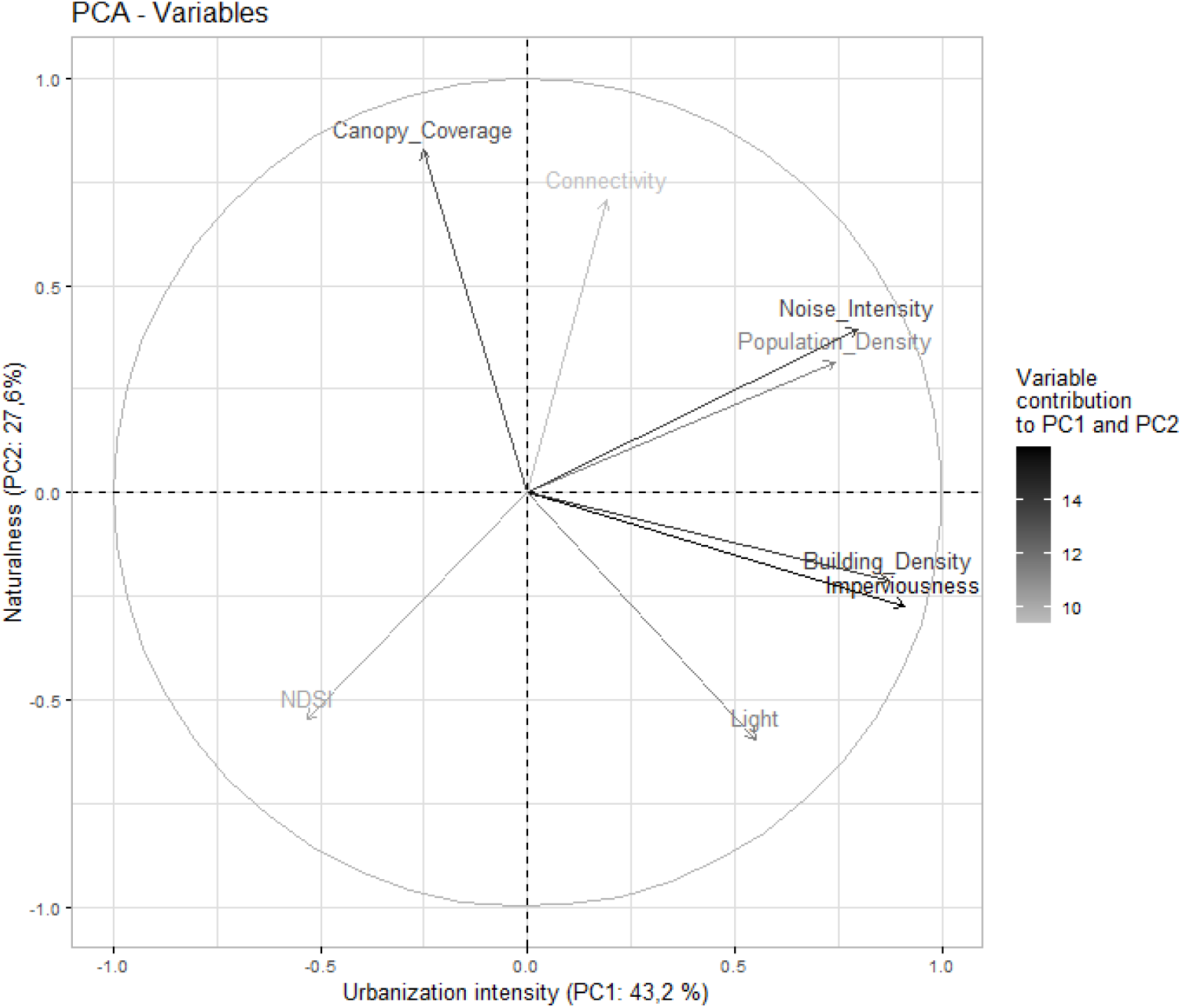
Variables factor map in Principal Components Analyses.

### 3.2. Bird species richness and functional diversity

We identified 43 bird species, among which 31 were classified as functional insectivores (Figure 3). The richness of functional insectivores varied from 2 to 16 species per urban plot (mean ± SD: 7.5 ± 3.7). The sum of occurrences of vocalizations per functional insectivore species varied from 1 to 142 across all sites (22.7 ± 37.7). However, across all sites, 85% of functional insectivore species emitted only one vocalization on average during ten minutes of recordings, and the number of vocalizations emitted at any given site never exceeded three within ten minutes. The American Robin, the House Sparrow, and the European Starling were the functional insectivore species with the most frequent vocalizations.

**Figure 3.**
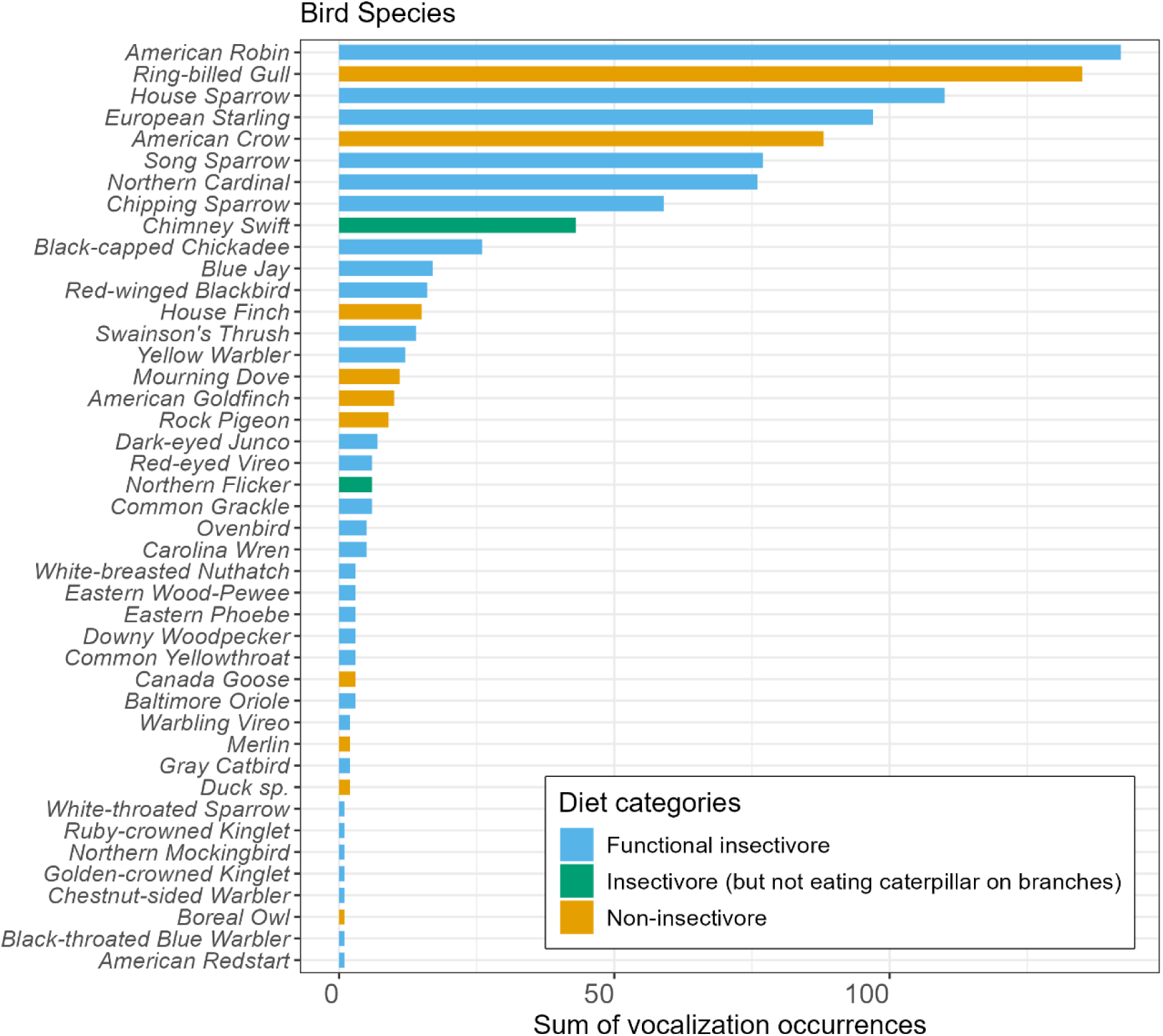
Decreasing total vocalization occurrences of species recorded, all urban plots combined. For this study, functional insectivorous species are the combination of the “strict insectivore” and “insectivore during the breeding season” categories.

Urbanization intensity (PC1) was the most important predictor for insectivore species richness (RVI=1.00), Shannon diversity (RVI=1.00), FDis (RVI=1.00), and BI (RVI=1.00).

Insectivore species richness, Shannon diversity and FDis consistently decreased whereas BI increased with increasing urbanization intensity (Figure 4). BI was strongly (RVI=1.00) and negatively (coefficient ± SE: -0.1777 ± 0.0640) influenced by tree species richness.

**Figure 4.**
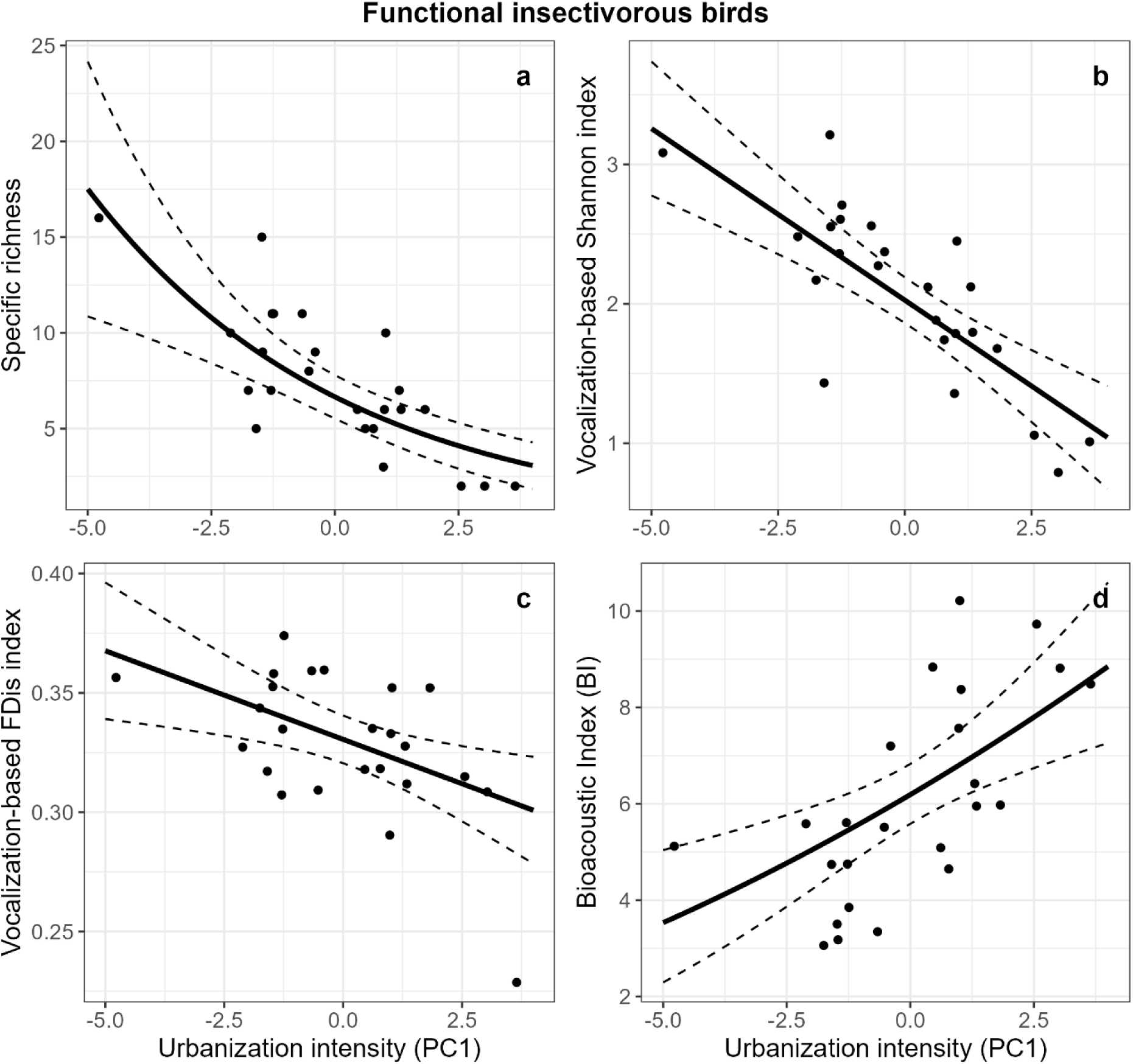
Statistical regression models explaining the relationships between (a) functional insectivore richness, (b) Shannon diversity, (c) FDis, (d) Bioacoustic Index and urbanization intensity (PC1). (a) is derived from a Poisson regression, (b) and (c) from linear regressions, and (d) from linear regression with square root transformation. Raw data are indicated by dots. The solid line represents the prediction from the models, dashed lines being the corresponding confidence intervals.

Insectivore species richness and Shannon diversity tended to be higher, and bird FDis tended to be lower in urban plots with greater tree specific richness. However, this predictor was considered less important than urbanization intensity for the three metrics (RVI=0.4, RVI=0.4 and RVI=0.32 for richness, Shannon diversity and FDis, respectively). FDis tended to positively respond to naturalness, but this predictor was of limited importance (RVI=0.59).

### 3.3. Bird predation attempts and herbivory regulation service

Of the 1,940 exposed plasticine caterpillars, 3.0 % (n = 59) had bird bill marks. Model selection retained nine models in the set of competing models in a range of ΔAICc < 2. Of the variables included in the set of competing models, tree species richness was the only significant one and was positively correlated with predation attempts (RVI = 1.00; see Fig.5). RVI of other predictors were all lower than 0.50 (Table SB).

**Figure 5.**
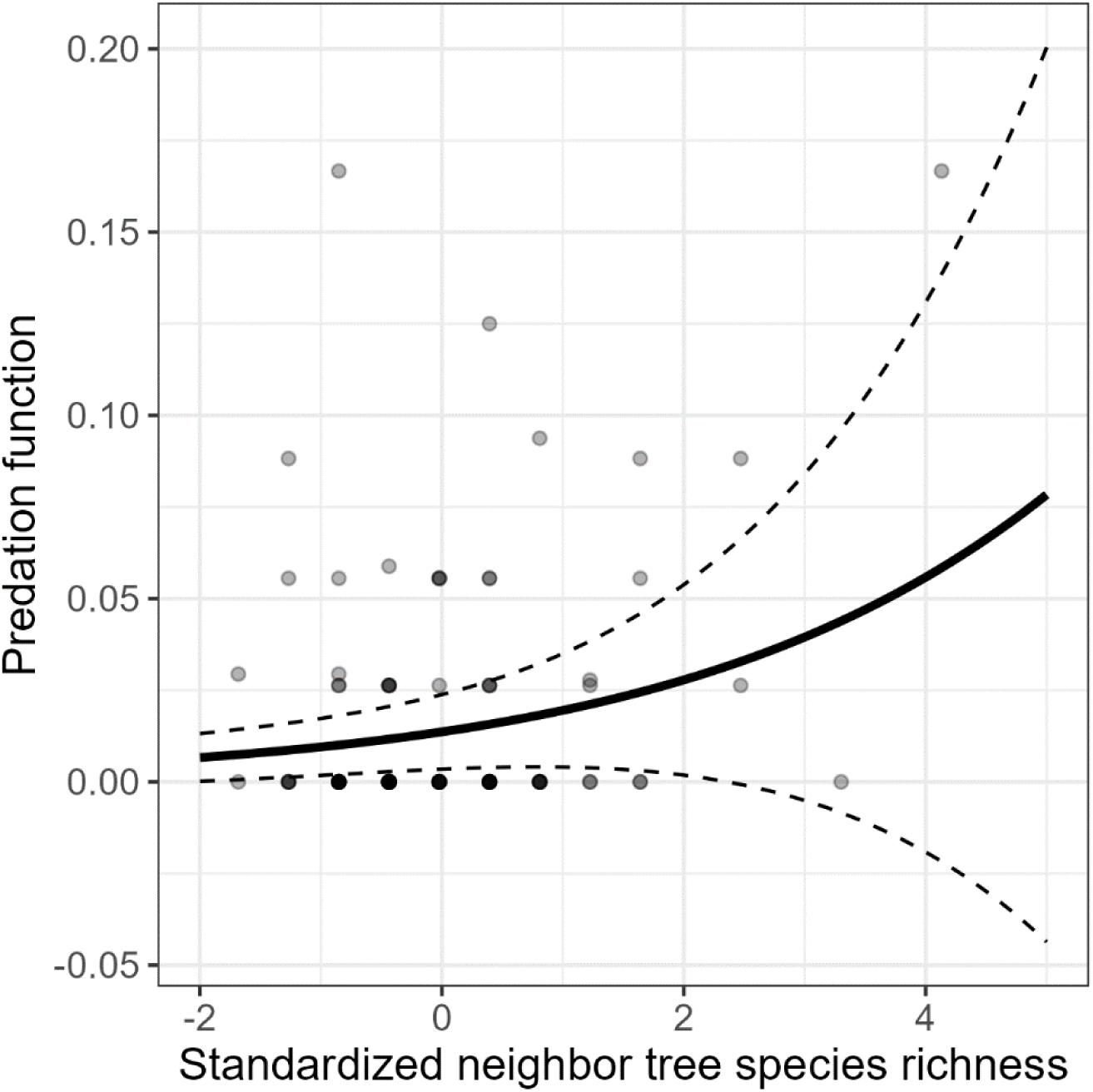
Relationship between bird predation attempts on artificial prey and tree species richness. Raw data are indicated by dots. Solid and dashed lines represent model prediction and corresponding confidence intervals. Grey intensity corresponds to the number of overlaying dots.

Insect herbivory was on average 0.09% higher in branches with bird exclusion than in control branches. These results are due to the very high variability in herbivory rates between tree species.

Urbanization intensity (RVI=1.00), tree species richness (RVI=1.00), and their interaction (RVI=1.00) were selected as important variables explaining variability in the herbivory regulation function provided by birds. The effect of urbanization intensity *per se* was however contingent on tree species richness (two-way interaction: -0.1847 ± 0.0613). Specifically, the regulating service of herbivory significantly decreases with tree species richness, but this relationship becomes less positive when urbanization intensity is very low. Bird species richness was retained in the second-best model, showing a positive correlation with the herbivory regulation function but with lower importance (RVI=0.32) (Fig.6, Table SB).

**Figure 6.**
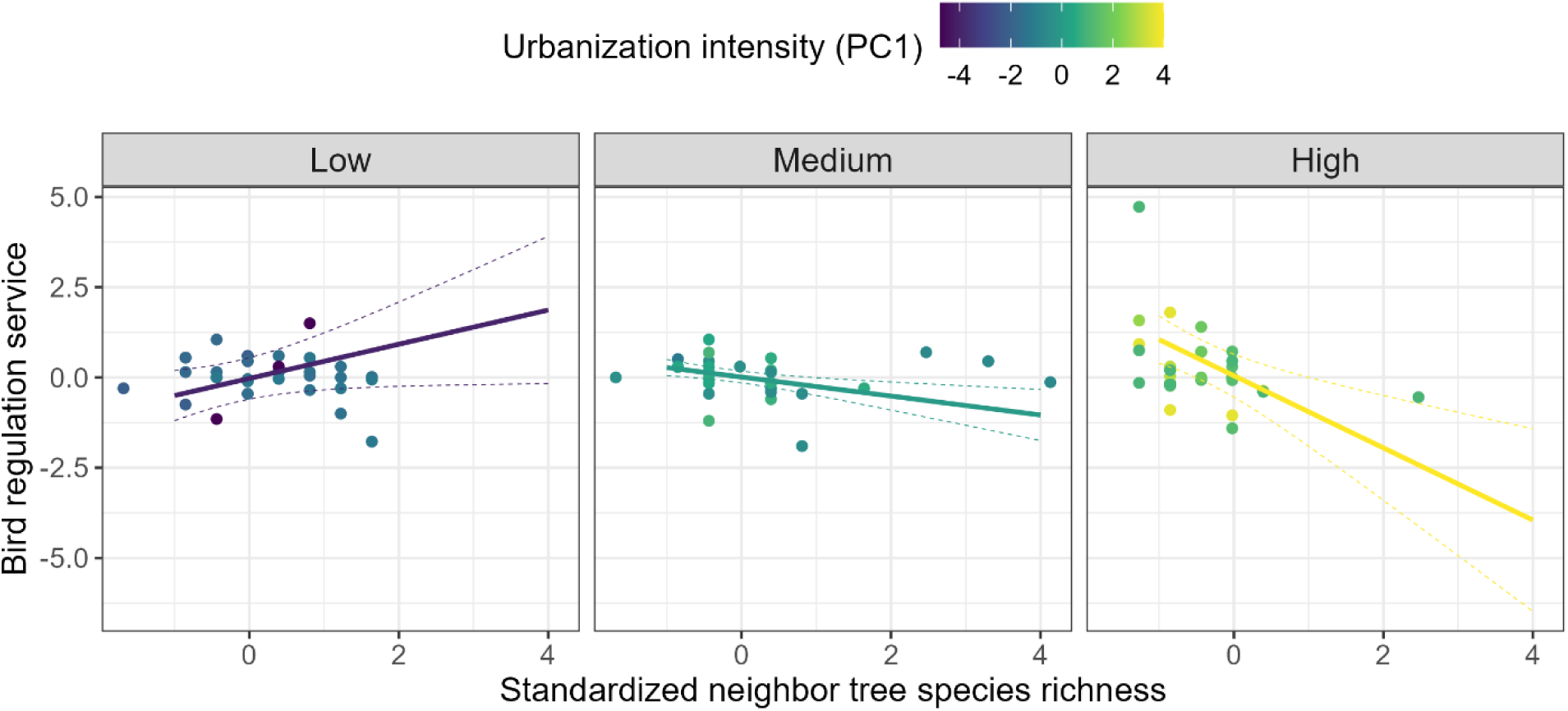
Regression-based on model predictions of changes in herbivory regulation function as a function of PC1 and neighbor tree species richness. Raw data are indicated by dots. The dashed lines represent the confidence intervals on the predictions.

## 4. Discussion

Our study unravels the complexity of the effect of urbanization on bird communities and the functions they support along an urbanization gradient. In particular, we found that the distribution and diversity of trees within the city modified the relationship between urbanization and bird insectivory, with important consequences for trophic interactions and ecological functions.

### 4.1. Urbanization influenced bird diversity, but not avian predation

Insectivorous bird species richness and vocalization-based functional diversity decreased with increasing urbanization within the boundaries of the city of Montreal. These findings are consistent with the long-standing theory that urbanization causes a reduction in the complexity of ecological communities (McKinney, 2006) as well as functional homogenization with more generalist species, which display greater resilience to human disturbances (Hahs et al., 2023). Research focusing on birds confirmed this general trend, both locally (Marcacci et al., 2021; Palacio et al., 2018; Santos et al., 2024) and globally (Sol et al., 2020). However, our study is one of the few (Carvajal-Castro et al., 2019; Yang et al., 2020) that have addressed the effect of within-city variability in insectivorous bird diversity in response to local factors related to the intensity of urbanization.

Previous studies have shown that avian insectivory could increase with the functional diversity of insectivorous birds (Barbaro et al., 2014; Philpott et al., 2009). Because insectivorous bird diversity decreased with increasing urbanization intensity, we expected that predation attempts on artificial prey as well as the effect of insectivorous birds on insect herbivory would also decrease along the urbanization gradient. Yet, we found no statistically significant relationship neither between urbanization and predation attempts nor between insectivorous bird diversity and predation attempts. Although literature reviews have questioned the generality of the relationship between urbanization and predation function (see Eötvös et al., 2018 for a meta-analysis), it is important to note that the majority of studies having addressed the effect of urbanization on tri-trophic interactions focused on comparisons between rural and urban habitats (Kozlov et al., 2017; Turrini et al., 2016) or on gradients spanning from these two extremes (Marcacci et al., 2021). The present study differs in that we addressed the effect of within-city variability in urbanization intensity. Still, in a rare comparable study, Long & Frank, (2020) reported higher avian predation attempts on plasticine caterpillars in urban forest remnants as compared to street trees.

The attack rate on artificial prey that we observed (3%) was lower than that reported in previous studies using the same methodology (*ca* 20%, e.g., Schillé et al., 2024; Valdés-Correcher et al., 2021), and even much lower than studies conducted in urban environments where it ranged between 30 and 70% (Kozlov et al., 2017; Long & Frank, 2020). In the context of our within-city study, we believe these low rates could be attributed to human disturbances at the study sites (often near houses or roads), increased caution of birds towards artificial prey in environments where they are frequently exposed to human waste that may resemble prey or even seasonal variability.

Some authors argue that predation attempts on artificial prey might be a poorly reliable proxy of predation (Zvereva & Kozlov, 2023). However, our conclusions remain the same regardless of whether we consider predation through the proportion of artificial prey with bill marks left by birds, or through the effect of bird exclusion on insect herbivory. We are therefore confident that the absence of a clear relationship between urbanization and predation on the one hand, and between bird diversity and predation on the other hand is not artefactual and reflects ecological features we did not capture. This is all the more true given that we did not observe a significant increase in herbivory between control branches and branches from which birds were excluded.

A possible explanation for the lack of correlation between urbanization or insectivorous bird diversity and predation rates might be that certain abundant, highly generalist bird species, such as House Sparrows and European Starlings, were the most likely agents attempting to prey upon plasticine prey in Montreal, particularly in highly urbanized areas. This explanation would be consistent with the high values of the Bioacoustic Index, which is a clue of high abundance and intense activity of particularly loud species (Boelman et al., 2007), like synurbic species, in the most urbanized plots while other birds’ vocalizations were more subdued in other plots. Although the Bioacoustic Index is only an indirect proxy of bird abundance that should be treated with caution, our personal field observations align with this. Indeed, House Sparrows, known for their opportunistic behavior and ability to scavenge on human food scraps (MacGregor-Fors et al., 2020), were especially prevalent in downtown Montreal, where we made direct observations of predation attempts on artificial prey immediately after installing them in trees.

We propose that the relatively constant predation rate we observed resulted from different mechanisms at both ends of the urbanization gradient we explored. In highly urbanized habitats, the absence of real prey may have increased the likelihood that generalist birds preyed on plasticine caterpillars. Conversely, in less urbanized areas, the greater abundance and diversity of birds may have increased the probability of encounters with lures, despite a higher availability of real prey. We cannot exclude the possibility that local factors, such as the presence of nest boxes or human bird-feeding behaviors, also influenced the relationship between urbanization, bird diversity, and predation. Only by considering both bird abundance and the availability of real prey can we make further inferences on the mechanisms at play, beyond mere speculation.

### 4.2. Tree diversity influenced bird predation, but not bird diversity

Trees generally promote bird diversity in cities by providing habitats and food for a diverse range of bird species (Dyson, 2020; Ferenc et al., 2014; May-Uc et al., 2020). However, contrary to our predictions, we found no evidence that tree density, distribution, or species richness were important in predicting bird diversity. It is possible that the distribution of treeless green spaces (Schütz & Schulze, 2015) as well as the vertical stratification of the vegetation (Campos-Silva & Piratelli, 2021) played a substantial role in bird abundance and diversity, which future studies should aim to better capture.

Likewise, we found that the Bioacoustic Index correlated negatively with tree species richness. Yet, because it reflects the abundance of species that vocalize with high intensity (Boelman et al., 2007), we expected the opposite. Two explanations can be proposed. First, in plots with high tree richness, dense foliage may dampen the sound intensity (Fang & Ling, 2003) whereas birds might have been more likely to have vocalized closer to the microphone in areas where the few available trees were those we focused on in this study. It is worth noting that in urban centers, where street trees are often isolated, loud vocalizations of Sparrows or Starlings were frequently recorded, as previously explained. Second, since the Bioacoustic Index is not a direct measure of bird abundance, some other sounds may share frequencies similar to those of birds, which overlap and cannot be disentangled in the automated calculation of this index (Bradfer-Lawrence et al., 2019).

Avian predation rate increased with increasing tree diversity, without releasing herbivory pressure. This finding is in line with the natural enemies’ hypothesis that predicts an increase in predation with increasing plant diversity (Stemmelen et al., 2022), but also adds to the controversy regarding the relationship between predator abundance, activity, and influence on trophic cascades (Staab & Schuldt, 2020). Unlike predation attempts on artificial prey, the trophic cascade whereby birds protect trees against insect herbivores – which we assessed through the difference in herbivory between netted and control branches – decreased in intensity with increasing tree diversity. This can be explained by an indirect effect of birds on insect herbivores through their predation upon mesopredators such as spiders, true bugs, and other predaceous arthropods (the “mesopredator release effect” Soulé et al., 1988). Changes in differential insect herbivory between control and netted branches may reflect either changes in overall avian predation, changes in predation by mesopredators, or both. Because predation attempts by birds on plasticine caterpillars increased with increasing tree diversity, we expected a concomitant decrease in herbivory in control branches as compared to netted ones. Yet, we found the opposite. We suggest this was due to higher avian predation on predaceous arthropods in areas with greater tree diversity, where their abundance and richness is expected to be higher (Stemmelen et al., 2022; Vehviläinen et al., 2008). Although differential avian predation on herbaceous vs. predaceous arthropods remains to be explored, such an explanation is consistent with the findings by Long & Frank (2020), who demonstrated predation attempts by arthropods on artificial prey in downtown Raleigh, NC, USA, are 30% more frequent than predation attempts by birds, whereas predation attempts by birds were 20% more frequent than predation attempts by arthropods within forest fragments. Our study did not account for arthropod predators, which would freely move through the mesh that excluded avian predators. A full understanding of trophic cascades in cities would therefore require that future studies quantify the relative contribution of vertebrate and invertebrate predation.

We acknowledge that the inconsistency in predation rate assessment questions the use of artificial prey, whose reliability is controversial (Zvereva & Kozlov, 2023). Still, despite legitimate concerns regarding the accuracy of predation clue identification (Bateman et al., 2017; Valdés-Correcher et al., 2022), and their interpretation (Nimalrathna et al., 2023; Rodriguez-Campbell et al., 2023), it is so far the most practical and relevant approach to assess bird foraging activity (Ferrante et al., 2024; Schillé et al., 2024).

### 4.3. Urbanization and tree diversity interactively influenced the top-down trophic cascade

We found that the effect of tree diversity on the trophic cascade was contingent on urbanization (and *vice versa*). Specifically, herbivore regulation by insectivorous birds increased with increasing tree diversity in the least urbanized areas of the city of Montreal, but its effect was the opposite in more urbanized areas. A likely explanation is that the low bird diversity in these areas prevented the few species present from benefiting from tree diversity, as they may only use a limited range of tree species. In contrast, in less urbanized areas, a higher diversity of birds could take advantage of greater tree species richness to find food, thereby increasing trophic interactions (Catterall et al., 1989; Nell et al., 2018; Strohbach et al., 2013).

### 4.4. Conclusion and implications for future research

In this study, we untied the complex interactions between trees, insect herbivores, and insectivorous birds in contrasting urban environments. We revealed a mismatch between the effect of urbanization on bird diversity and its effect on the regulation service, therefore generating unusual diversity-function relationships. We also found that the intensity of the trophic cascade whereby insectivorous birds influence insect herbivory in trees was modulated by the diversity of trees at the local scale. Our results have several implications for further fundamental and applied research on the urban forest and associated biodiversity. First, we show that urban biodiversity as well as the ecological functions it supports cannot be fully understood without accounting for within-city heterogeneity in urbanization intensity.

We therefore call for a more specific description of urbanization in future ecological studies that better account for and disentangle the intertwined factors of urbanization. Second, because biodiversity is made of moving organisms, we emphasize the critical need to expand our knowledge of urban biodiversity beyond the limits of the public domain. This represents a major challenge for future ecological research in cities and for cities that will require strengthened collaborations between the public, researchers, and local authorities. Finally, we stressed the need to expand the study of urban biodiversity to its functional dimension in order to fully understand its implications for the functioning of urban ecosystems.

## Data availability statement

For the moment, there is an embargo on data and codes, which will be lifted after open acceptance.

## Copyright statement

For the purpose of Open Access, a CC-BY 4.0 public copyright license () has been applied by the authors to the present document and will be applied to all subsequent versions up to the Author Accepted Manuscript arising from this submission.

## Acknowledgement

LS was supported by grants from the Science and Environment Doctoral School of the University of Bordeaux, from the Graduate Schools of the University of Bordeaux, from the Agreenium International Research School (EIR-A) and from Erasmus+. This study was permitted by the financial support of the BNP Paribas Foundation through its Climate & Biodiversity Initiative for the ‘Tree bodyguards’ citizen science project.

## Author contribution

B.C., L.B., A.P., and L.S. conceptualized the study and developed the methodology. L.S., G.M., and S.C. collected the data. H.O. processed audio recordings for bird species identification. L.S. processed and analyzed the data with guidance from B.C. and L.B. L.S., B.C., L.B. led the writing and all authors contributed critically to the revisions.

