## Supplemental Table 1 and Table 2 for "Urban tree diversity fosters bird insectivory despite a loss in bird diversity with urbanization"

### Supplementary material

*Table SA. Identity of focal trees for each urban plot*

| Plot_ID | Tree_ID | Tree_species |
|---|---|---|
| 01A | 32 | *Acer saccharinum* |
| 01A | 91 | *Acer platanoides* |
| 01A | 107 | *Acer platanoides* |
| 02A | 188 | *Gleditsia triacanthos* |
| 02A | 195 | *Ulmus americana* |
| 02A | 286 | *Gleditsia triacanthos* |
| 02A | 252 | *Acer platanoides* |
| 03A | 292 | *Gleditsia triacanthos* |
| 03A | 299 | *Ulmus pumila* |
| 03A | 300 | *Acer saccharinum* |
| 03A | 354 | *Tilia cordata* |
| 04A | 522 | *Gleditsia triacanthos* |
| 04A | 530 | *Ulmus americana* |
| 04A | 525 | *Gleditsia triacanthos* |
| 04A | 520 | *Gleditsia triacanthos* |
| 05B | 753 | *Gleditsia triacanthos* |
| 05B | 762 | *Gleditsia triacanthos* |
| 05B | 765 | *Acer platanoides* |
| 05B | 773 | *Acer platanoides* |
| 05B | 775 | *Acer platanoides* |
| 06C | 888 | *Ulmus pumila* |
| 06C | 898 | *Acer saccharinum* |
| 06C | 887 | *Acer platanoides* |
| 06C | 893 | *Ulmus pumila* |
| 07A | 1010 | *Gleditsia triacanthos* |
| 07A | 1012 | *Gleditsia triacanthos* |

| | | |
|---|---|---|
| 07A | 1024 | *Acer platanoides* |
| 07A | 1019 | *Gleditsia triacanthos* |
| 08A | 1089 | *Gleditsia triacanthos* |
| 08A | 1090 | *Ulmus x 'Sapporo Autumn Gold'* |
| 08A | 1097 | *Tilia americana* |
| 08A | 1109 | *Tilia americana* |
| 09B | 1217 | *Prunus spp.* |
| 09B | 1233 | *Acer saccharinum* |
| 09B | 1241 | *Acer platanoides* |
| 09B | 1262 | *Acer platanoides* |
| 10A | 1366 | *Prunus spp.* |
| 10A | 1356 | *Gleditsia triacanthos* |
| 10A | 1398 | *Tilia americana* |
| 10A | 1407 | *Gleditsia triacanthos* |
| 11B | 1487 | *Gleditsia triacanthos* |
| 11B | 1481 | *Gleditsia triacanthos* |
| 11B | 1499 | *Ulmus x 'Homestead'* |
| 11B | 1522 | *Ulmus americana* |
| 12B | 1597 | *Ulmus x 'Homestead'* |
| 12B | 1624 | *Ulmus x 'Homestead'* |
| 12B | 1621 | *Acer saccharinum* |
| 12B | 1623 | *Gleditsia triacanthos* |
| 12C | 1689 | *Ulmus americana* |
| 12C | 1704 | *Ulmus americana* |
| 12C | 1710 | *Gleditsia triacanthos* |
| 12C | 1706 | *Ulmus x 'Morton'* |
| 13A | 1828 | *Acer saccharinum* |
| 13A | 1835 | *Acer negundo* |
| 13A | 1841 | *Acer platanoides* |
| 13A | 1869 | *Gleditsia triacanthos* |

| | | |
|---|---|---|
| 14A | 1936 | *Ulmus x 'Morton'* |
| 14A | 1986 | *Tilia americana* |
| 14A | 2028 | *Tilia cordata* |
| 14A | 2039 | *Tilia cordata* |
| 15B | 2186 | *Acer platanoides* |
| 15B | 2204 | *Gleditsia triacanthos* |
| 15B | 2206 | *Gleditsia triacanthos* |
| 15B | 2201 | *Ulmus americana* |
| 16A | 2347 | *Acer platanoides* |
| 16A | 2301 | *Acer platanoides* |
| 16A | 2362 | *Acer negundo* |
| 16A | 2445 | *Prunus spp.* |
| 17C | 2521 | *Acer platanoides* |
| 17C | 2551 | *Acer platanoides* |
| 17C | 2567 | *Gleditsia triacanthos* |
| 17C | 2571 | *Gleditsia triacanthos* |
| 18B | 2718 | *Acer saccharinum* |
| 18B | B | *Ulmus x 'Morton'* |
| 18B | 2644 | *Acer saccharinum* |
| 19B | 2743 | *Acer negundo* |
| 19B | C | *Acer negundo* |
| 19B | 2760 | *Tilia cordata* |
| 19B | 2778 | *Acer saccharinum* |
| 21B | 2823 | *Ulmus pumila* |
| 21B | 2856 | *Gleditsia triacanthos* |
| 21B | 2868 | *Acer negundo* |
| 21B | 2864 | *Acer negundo* |
| 22A | 2980 | *Gleditsia triacanthos* |
| 22A | 2983 | *Acer saccharinum* |
| 22A | 2990 | *Tilia americana* |

| | | |
|---|---|---|
| 22A | 3016 | *Tilia americana* |
| 23A | 3103 | *Acer platanoides* |
| 23A | 3123 | *Acer platanoides* |
| 23A | 3093 | *Acer platanoides* |
| 23A | 3132 | *Acer platanoides* |
| 24B | 3188 | *Acer platanoides* |
| 24B | 3200 | *Ulmus x 'Patriots'* |
| 24B | 3208 | *Acer platanoides* |
| 24B | 3228 | *Acer saccharinum* |

*Table SB. Summary of model selection for each of the three sets of linear models used to analyze (a) taxonomic or functional diversity (any of the three indices); (b) bird predation attempts and (c) herbivory. Explanatory variables that cannot be included in the model selection for a given response variable are shaded. Values correspond to model coefficients for variables included in a particular model. Values in bold indicate that the coefficient was significantly different from zero. The gray line with white font following each model selection displays the relative variable importance (RVI) of each selected variable in the model selection process (i.e., the sum of weights from each model including that variable).*

*df : degrees of freedom ; logLik : model log-likelihood ; AICc : Akaike information criterion adapted to small sample sizes ; ΔAICc : difference between model AICc and that of the best model ; only models with ΔAICc < 2 are shown ; R2 conditional and marginal are also shown for mixed models.*

| | Response variable | PC1 | PC2 | Tree Specific richness | Dim.1 : Tree Specific richness | Dim.2 : Tree Specific richness | Specific Richness of Funct Insect | BI | df | logLik | AICc | ΔAICc | weight | $R^2_{marginal}$ | $R^2_{conditionnal}$ |
|---|---|---|---|---|---|---|---|---|---|---|---|---|---|---|---|
| | Specific Richness of Funct Insect | **-0.2002** | | | | | | | 2 | -53.815 | 112.2 | 0.00 | 0.418 | 0.59 | |
| | Specific Richness of Funct Insect | **-0.1933** | | 0.09739 | | | | | 3 | -52.907 | 113.0 | 0.81 | 0.279 | 0.63 | |
| | **RVI** | **1.00** | | 0.4 | | | | | | | | | | | |
| | Vocalization-based Shannon index | **-0.2647** | | | | | | | 3 | -10.566 | 28.3 | 0.00 | 0.451 | 0.62 | |
| a | Vocalization -based Shannon index | **-0.2462** | | 0.11630 | | | | | 4 | -9.522 | 29.1 | 0.82 | 0.300 | 0.64 | |
| | **RVI** | **1.00** | | 0.4 | | | | | | | | | | | |
| | Vocalization -based FDis | **-0.008857** | | | | | | | 3 | 54.815 | -102.4 | 0.00 | 0.294 | 0.30 | |
| | Vocalization -based FDis | **-0.007425** | 1.134e-03 | -0.002179 | | **-0.01149** | | | 6 | 59.436 | -101.9 | 0.50 | 0.229 | 0.48 | |
| | Vocalization -based FDis | **-0.008857** | 4.686e-03 | | | | | | 4 | 55.814 | -101.5 | 0.91 | 0.187 | 0.34 | |
| | **RVI** | **1.00** | 0.59 | 0.32 | | **0.32** | | | | | | | | | |
| | Bioacoustic Index (BI) | **0.12170** | | **-0.1777** | | | | | 4 | -2.945 | 16.0 | 0.00 | 0.586 | 0.56 | |
| | **RVI** | **1.00** | | **1.00** | | | | | | | | | | | |
| | Bird predation attempts | | | **0.3156** | | | | | 4 | -98.075 | 204.6 | 0.00 | 0.105 | 0.33 | 0.87 |
| | Bird predation attempts | 0.05565 | 0.19100 | **0.2726** | -0.3126 | | | | 7 | -94.867 | 205.0 | 0.42 | 0.085 | 0.33 | 0.86 |
| | Bird predation attempts | 0.06407 | | **0.2857** | -0.2055 | | | | 6 | -96.067 | 205.1 | 0.49 | 0.082 | 0.33 | 0.87 |
| | Bird predation attempts | 0.05748 | 0.19430 | **0.3489** | -0.2860 | 0.1234 | | | 8 | -93.942 | 205.5 | 0.95 | 0.065 | 0.32 | 0.86 |
| b | Bird predation attempts | 0.11080 | | **0.3645** | | | | | 5 | -97.529 | 205.7 | 1.14 | 0.059 | 0.33 | 0.87 |
| | Bird predation attempts | | 0.08911 | **0.4061** | | 0.1854 | | | 6 | -96.482 | 205.9 | 1.32 | 0.054 | 0.34 | 0.87 |
| | Bird predation attempts | | | **0.3675** | | | -0.166300 | | 5 | -97.692 | 206.1 | 1.46 | 0.050 | 0.34 | 0.87 |
| | Bird predation attempts | | 0.07120 | **0.3208** | | | | | 5 | -97.906 | 206.5 | 1.89 | 0.041 | 0.32 | 0.86 |
| | Bird predation attempts | | | **0.3511** | | | | 0.09801 | 5 | -97.942 | 206.6 | 1.96 | 0.039 | 0.33 | 0.86 |
| | **RVI** | 0.50 | 0.42 | **1.00** | 0.40 | 0.21 | 0.09 | 0.07 | | | | | | | |
| | Difference in herbivory | -0.035890 | | **-0.2524** | **-0.1869** | | | | 7 | -104.291 | 223.9 | 0.00 | 0.264 | 0.14 | 0.14 |
| c | Difference in herbivory | 0.012200 | | **-0.2646** | **-0.1800** | | 0.11750 | | 8 | -103.858 | 225.4 | 1.53 | 0.123 | 0.15 | 0.15 |
| | **RVI** | 1.00 | | **1.00** | **1.00** | | 0.32 | | | | | | | | |
